# GPR171 Agonist Reduces Chronic Neuropathic and Inflammatory Pain in Male, but not in Female Mice

**DOI:** 10.1101/2021.04.16.440030

**Authors:** Akila Ram, Taylor Edwards, Ashley McCarty, Leela Afrose, Max V. McDermott, Erin N. Bobeck

## Abstract

Chronic pain is a growing public health crisis that requires exigent and efficacious therapeutics. GPR171 is a promising therapeutic target that is widely expressed through the brain, including within the descending pain modulatory regions. Here, we explore the therapeutic potential of the GPR171 agonist, MS15203, in its ability to alleviate chronic pain in male and female mice using a once-daily systemic dose (10mg/kg, i.p.) of MS15203 over the course of 5 days. We found that in our models of Complete Freund’s Adjuvant (CFA)-induced inflammatory pain and chemotherapy-induced peripheral neuropathy (CIPN), MS15203 did not reduce thermal hypersensitivity and allodynia, respectively, in female mice. On the other hand, MS15203 treatment decreased the duration of thermal hyper-sensitivity in CFA-treated male mice following 3 days of once-daily administration. MS15203 treatment also produced an improvement in allodynia in male mice, but not female mice, in neuropathic pain after 5 days of treatment. Gene expression of GPR171 and that of its endogenous ligand BigLEN, encoded by the gene PCSK1N, were unaltered within the periaqueductal gray in both male and female mice following inflammatory and neuropathic pain. However, following neuropathic pain in male mice, the protein levels of GPR171 were decreased in the periaqueductal gray. Treatment with MS15203 then rescued the protein levels of GPR171 in the periaqueductal gray of these mice. Taken together, our results identify GPR171 as a GPCR that displays sexual dimorphism in alleviation of chronic pain. Further, our results suggest that GPR171 and MS15203 have demonstrable therapeutic potential in the treatment of chronic pain.

## Introduction

Chronic pain is growing public health concern with over 50 million adults in the United States having suffered from the condition since 2016 (1). Globally, nearly 2 billion people have been affected by chronic pain conditions, emphasizing the need for extensive research and development of efficacious therapeutics (2). Despite the progress made in studying the causal and therapeutic mechanisms of chronic pain, the inclusion of female cohorts in pain studies is limited (3, 4). Several reports have highlighted the existence of sex differences in the pathology of chronic pain (5–8). Indeed, females have been reported to experience more severe and persistent pain than males in postoperative settings leading to reduced levels of physical activity (9). In addition, the estrus stage of females has also been implicated in their pain sensitivity and their responsiveness to opioid analgesics (10, 11). However, the long-term use of opioid analgesics for the treatment of chronic pain has profound negative side effects and has been shown to have limited effectiveness in the daily management of chronic pain (12). Given these substantial issues with usage of opioids, we sought to look beyond this class of drugs to identify efficacious therapeutics for chronic pain.

G-protein coupled receptors (GPCRs) and their dysregulation is implicated in a wide range of pathologies, including chronic pain (13–15). However, only 34% of FDA-approved drugs target GPCRs, highlighting the yet untapped potential of the GPCR family to be therapeutic targets (16, 17). One particularly promising target of interest for novel pain therapeutics is the G-protein coupled receptor, GPR171, and its endogenous ligand, BigLEN. The peptide BigLEN is derived from the neuropeptide precursor ProSAAS. The precursor peptide, in turn, has been shown to be upregulated in the cerebrospinal fluid of patients with fibromyalgia and within the periaqueductal gray in a rodent model of opioid-induced hyperalgesia (18, 19). The receptor, GPR171, and its endogenous ligand, BigLEN, are widely expressed through the brain including the periaqueductal gray (PAG) (20, 21). This particular brain region is situated in the descending pain modulatory pathway and is a site of action for a range of antinociceptive drugs including opioids and cannabinoids (22, 23). Previous studies exploring the role of GPR171 in acute pain showed that agonism of the receptor via systemic administration of the synthetic agonist, compound MS15203, led to an increase in the antinociceptive effect of systemic morphine administration (20).

We explored the role of GPR171 in chronic neuropathic and inflammatory pain in male and female mice. The therapeutic potential of the GPR171 agonist compound, MS15203, was assessed via repeated once-daily systemic (10 mg/kg i.p.) injections. We found that the compound MS15203 reduced the duration of chronic neuropathic and inflammatory pain in male mice, but not female mice. While we found no alterations in gene expression levels of ProSAAS or GPR171 in the PAG of male or female mice, we found that CIPN produces a decrease in GPR171 protein levels in vlPAG of male mice. We note that following MS15203 treatment, the GPR171 protein levels in male mice with neuropathic pain recovered and were indeed elevated compared to untreated controls. Our findings demonstrate a sexually dimorphic receptor system in chronic pain and establish a role for the recently deorphanized receptor GPR171 in the reduction of chronic pain in male mice.

## Materials and Methods

### A. Animals

102 Adult male and female C57BL/6CS mice (Charles River Laboratories, CA), 6-8 weeks old and weighing 18-26 g at the start of the study were used in the study. Food and water were available ad libitum, except during testing. Mice were housed (four to five per cage) in a humidity and temperature-controlled room with a 12-hour light/dark cycle (off at 1900). All behavior testing took place during the light cycle. For the female cohorts, estrus stage was determined by obtaining a vaginal lavage daily starting 5 days prior to the start of the study. A lavage was also obtained daily over the course of the study. All procedures were performed in accordance with the Guide for the Care and Use of Laboratory Animals adopted by the National Institutes of Health and were approved by the Institutional Animal Care and Use Committee of Utah State University (Protocol 2775).

### B. CFA-induced inflammatory pain

Animals were very briefly anesthetized with isofluorane (99.9%, inhaled; Fluriso, VetOne) and restrained in order to access the plantar surface of the hind paw. Complete Freund’s Adjuvant (CFA, Cat. F5881, Sigma-Aldrich) was injected under the epidermis into the plantar surface of both hind paws using a 27G needle (20 *µ*l/paw) as described previously (24). Animals in the control group received light isofluorane anesthesia alone without manipulation of their hind paws to prevent inflammation from the injection procedure. Drug administration commenced 24h following CFA injection.

### C. Assessment of inflammatory pain in mice

A plantar test was performed as previously described (25) to assess inflammatory pain. Briefly, the subjects were placed in plexiglass enclosures on top of a glass platform. The animals were acclimatized to the testing chamber for 3 days prior to the start of the study (1 hour/session). On each testing day, the animals were acclimatized in the testing chamber for 20 minutes, or until cessation of exploratory behavior. A radiant heat source (IITC, Cat. 390) was applied to the plantar surface of the hind paw and the time to a nocifensive response was recorded. A cut-off time of 20 seconds was enforced to avoid potential injury due to tissue damage. Two trials were performed on each hindpaw to obtain the average reaction time per paw and a third reaction time was obtained if the preceding two values differed by 2s or more. Both left and right hindpaws were tested and their average thermal latency was considered for statistical analysis. Baseline latencies were established prior to injection of CFA. Following CFA injection on Day 0, the plantar test took place on Days 1, 3, and 5, where Day 0 marks the injection of CFA and Day 1 is 24h following CFA injection.

### D. Chemotherapy-induced peripheral neuropathy

Paclitaxel (Sigma-Aldrich, MO, USA) was diluted in a vehicle comprising of Cremphor (Sigma-Aldrich, MO, USA)/90% ethanol/ 0.9% saline in a 1:1:18 ratio, and administered intraperitoneally (i.p.; 4 mg/kg) to mice on four alternate days (cumulative dose 16 mg/kg) to induce neuropathy as described previously (26, 27). The control group received four injections of the vehicle (10 mL/kg). As paclitaxel can be present in animal excreta, bedding of mice used in this study was treated as a biohazard and disposed according to University guidelines.

### E. Assessment of mechanical allodynia in mice

The von Frey filament test was performed as previously described (26) to assess allodynia in mice. Briefly, all mice (vehicle- and paclitaxel-treated) were placed in plexiglass enclosures mounted onto a testing platform containing a metal, perforated floor (Stoelting Co., Wood Dale, IL, USA). Animals were acclimatized to the testing chamber for 3 days prior to the start of the study (1 hour/session). On each testing day, the animals were acclimatized in the testing chamber for 20 minutes, or until cessation of exploratory behavior. Mechanical allodynia was assessed in male mice by applying von Frey filaments to the midplantar region of both hindpaws for approximately 2s per stimulus using calibrated filaments (Touch Test kit, Cat. NC12775-99, North Coast Medical). All trials began with the 1g filament and proceeded using an up-down trial design. Both right and left hindpaws were tested and their average mechanical threshold was considered for statistical analysis. A sudden paw withdrawal, flinching, or paw licking was regarded as a nocifensive response. A negative response was followed by the use of a larger filament. For assessment of mechanical allodynia in females, an electronic von Frey device (Ugo Basile, Italy; Cat. 38450) was used as the allodynic females’ mechanical thresholds were lower than the detectable range of the manual filaments. Paw mechanical withdrawal thresholds are expressed as a percentage of baseline values, where the baseline represents von Frey thresholds prior to paclitaxel or vehicle treatment.

### F. Drug treatment

GPR171 agonist (MS15203, a gift from Dr. Sanjai Pathak) was dissolved in sterile 0.9% saline (1 mg/ml). Mice in the Paclitaxel- and CFA-induced chronic pain studies were randomly assigned, by block randomization, into four treatment groups: pain + MS15203, pain + saline, no pain + MS15203, and no pain + saline. Mice in the MS15203 treatment group received 10 mg/kg i.p. as this dose was previously shown to produce an increase in hot plate thermal latency when co-administered with morphine (20). To evaluate the effect of MS15203 on chronic neuropathic pain, the animals were administered with one dose of either MS15203 or saline, once daily for 5 days starting on Day 15. To evaluate the effect of MS15203 on chronic inflammatory pain, the animals were administered with one dose of either MS15203 or saline, once daily for 5 days starting 24h following induction of inflammatory pain.

### G. Immunofluorescence staining and microscopy

Immediately following the final behavioral testing, a subset of the subjects (3/group) were deeply anesthetized using isofluorane and transcardially perfused with 4% paraformaldehyde (PFA). Their brains were post-fixed in 4% PFA for 1 hour and then were stored in 1x PBS at 4°C until further processing. Sectioning was performed on a vibratome (Leica Biosystems, Germany) and 50 *µ*m sections containing the vlPAG were selected for immunofluorescence analysis. The sections were first incubated in sodium borohydride (1% in PBS) to expose the epitopes and decrease autofluorescence from aldehydes. Subsequently, they were permeabilized with Triton-X 100 (3%, Sigma-Aldrich), blocked with normal goat serum (5% in PBS), and incubated overnight in primary antibodies in their appropriate buffer (1% BSA (Sigma-Aldrich) in PBS). The following day, the slices were washed in PBS and incubated for 2 hours in diluted secondary antibodies in 1% BSA in PBS. The antibodies were sourced from GeneTex (Rabbit anti-GPR171, Cat. GTX108131, 1:400) and Life Technologies (Goat anti-Rabbit A594, Cat. A11037, 1:1000). Following subsequent PBS washes, the slices were briefly (5 mins) incubated in DAPI, washed with PBS and mounted with an anti-fade (ProLong Diamond Anti-Fade, Cat. P36961, Invitrogen) on a glass slide. Representative images of coronal sections containing the vlPAG, as defined by the mouse brain atlas (28), were acquired on a Zeiss LSM 710 confocal microscope (Carl Zeiss Microscopy, Germany) at Utah State University’s microscopy core facility.

### H. Quantification of immunofluorescence

Quantification of GPR171-fluorescence was performed using ImageJ (NIH) on automatically thresholded images as previously described (29, 30). Up to 5 images of the vlPAG representative of both left and right hemispheres and across different rostrocaudal positions were analyzed per animal per group. They were analyzed as independent measurements to account for the structural and functional heterogeneity of the PAG (31– 34). Ten cells in the field of view were randomly selected along with three areas representing background fluorescence. The fluorescence from GPR171 signal was calculated using the formula for Corrected Total Cell Fluorescence (CTCF) (Integrated Density – (area of selection×mean background fluorescence)). The measurements have been presented in arbitrary units (AU).

### I. Quantitative RT-PCR

Immediately following the final behavioral testing, a subset of the subjects (3-5/group) were euthanized by decapitation and the periaqueductal gray was dissected and snap-frozen on dry ice. The tissue samples were stored at -80°C until further processing. RNA was extracted from the tissues using Trizol (Cat. 15596026, Invitrogen) and RNeasy Plus Mini Kit (Cat. 74136, Qiagen) following the manufacturer’s instructions. The eluted RNA was quantified using a Qubit 4 fluorometer (Invitrogen). cDNA was prepared using the Maxima first-strand synthesis kit for qRT-PCT (Cat. K1642, Thermo-Fisher) following the manufacturer’s instructions. Samples were prepared using SYBR green (iTaq Universal SYBR Green Supermix, Cat. 1725121, Bio-Rad, CA) for gene expression analysis on a real-time thermocycler (CFX384 Touch, BioRad). Custom primers as described previously (35) were used for GAPDH, GPR171, and ProSAAS (Integrated DNA Technologies). The synthesized cDNA was assayed in triplicate and analyzed using the 2△△Ct method. Here, Ct indicates the cycle number at which the fluorescence signal crosses an arbitrary threshold within the exponential phase of the amplification curve. To calculate △△Ct, we used the formula △△Ct = {(Cttarget:treated sample – Ct_GAPDH:treated sample_) – (Ct_target:control sample_ – Ct_GAPDH:control sample_)}. The value of the control sample was set to 100% and all samples were evaluated with respect to the control. Negative control reactions were performed to ascertain contaminant-free cDNA synthesis and primer specificity was evaluated using melt curve analyses. The primer sequence information can be found in the Supplementary Material (Table S1).

### J. Statistical Analysis

Results are presented as mean ± standard error of mean (SEM). Statistical analyses of behavioral studies were performed using a repeated measures two way analysis of variance (ANOVA) combined with Bonferroni’s posthoc test. Normality of data was evaluated using the D-Agostino and Pearson test. Statistical analyses of qRT-PCR data and fluorescence quantification were performed using a one-way ANOVA combined with a Tukey’s posthoc test. Experimenters were blinded to drug treatment (saline and MS15203), but not to the pain state during behavioral studies and statistical analysis. All graphing and statistical analyses were performed using Prism 9.0 (GraphPad, San Diego, CA).

## Results

### A. GPR171 agonist does not reduce chronic inflammatory pain in female mice

We used a model of CFA-induced inflammatory pain to study the effect of GPR171 agonism on pain relief (Fig. 1A). In females, (n=6-8/group), intra-plantar CFA injection produced inflammatory pain and associated thermal hypersensitivity as measured by a plantar test. The estrus stage did not impact the subjects’ thermal thresholds (Fig. S1A). A repeated measures two-way ANOVA indicated significant differences in thermal thresholds over the course of the study (Time [F (3.159, 88.44) = 36.78, p<0.0001]; Treatment [F (3, 28) = 14.57, p<0.0001]; Time × Treatment [F (12, 112) = 8.211, p<0.0001]). A Bonferroni’s post-hoc test revealed that the female mice developed thermal hypersensitivity 24 hours following the CFA insult (Fig. 1B: Day 1, Control + Saline vs CFA + Saline or CFA + MS15203, p<0.001) which persisted through the duration of the study (Fig. 1B: Day 5 post-drug, Control + Saline vs CFA + Saline, p<0.0001). However, the Bonferroni’s post-hoc analysis revealed that despite chronic (once-daily) treatment with MS15203, there was no change in the thermal latency of female mice with inflammation-induced hypersensitivity (Fig. 1B: Day 5 post-drug, CFA + Saline vs CFA + MS15203, p>0.05). MS15203 treatment alone did not alter the thermal latencies of females in the Control group.

**Fig. 1.**
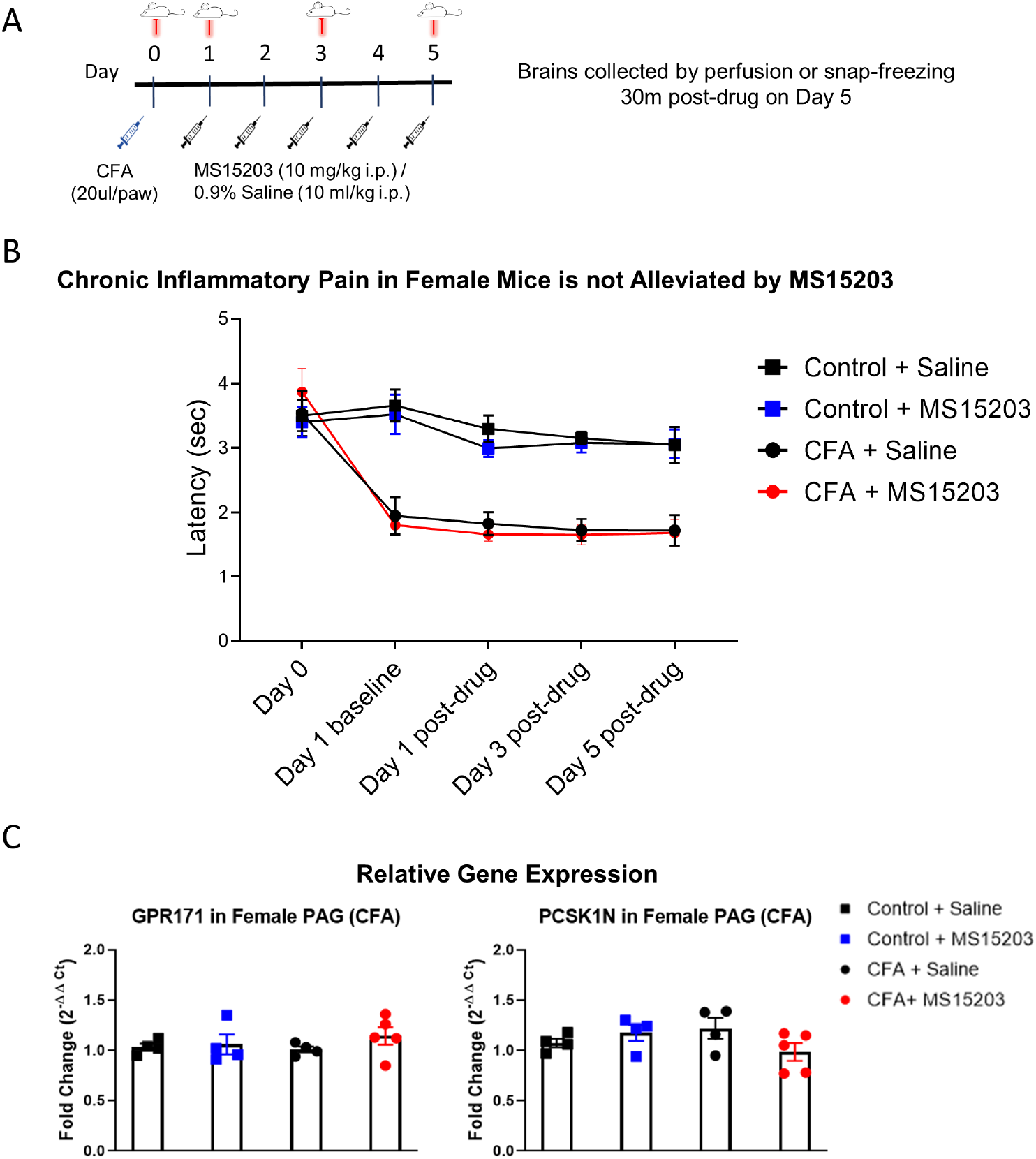
GPR171 agonist does not reduce chronic inflammatory pain in female mice. (A) An illustrative timeline describing the experimental design of CFA-induced inflammatory pain followed by MS15203 treatment. (B) Female mice (n=6-8/group) injected with CFA in their hind paws developed rapid thermal hypersensitivity as measured by a plantar test. Chronic treatment with MS15203 failed to reduce inflammatory pain as noted by the persistence of thermal hypersensitivity through Day 5. Repeated measured two-way ANOVA with Bonferroni’s post-hoc test. (C) Analysis of gene expression changes within the periaqueductal gray (PAG) reveals that the transcript levels of GPR171 and its endogenous ligand, PCSK1N, are unchanged in female mice (n=4-5/group) irrespective of inflammatory pain status or MS15203 treatment. One-way ANOVA with Tukey’s post-hoc test.

We then performed qRT-PCR on dissected whole periaqueductal gray (PAG) tissue from a subset of females (n=4-5/group) to evaluate gene expression changes in GPR171 or its endogenous ligand, PCSK1N (Fig. 1C). The PAG is a critical modulator of antinociception within the descending pain pathway and is a site of action of endogenous opioid activity (32, 36). A one-way ANOVA indicated no significant differences between treatment groups on evaluation of GPR171 (Treatment F (3, 13) = 0.71, p>0.05) or PCSK1N (Treatment F (3, 13) = 1.68, p>0.05) expression in the PAG.

### B. GPR171 agonist reduces chronic inflammatory pain in male mice

We performed a similar experimental paradigm as Fig. 1A to evaluate the analgesic properties of GPR171 in male mice. In males, (n=5-6/group), intraplantar CFA injection produced inflammatory pain and thermal hypersensitivity on the plantar test. A repeated measures two-way ANOVA indicated significant differences in thermal thresholds over the course of the study (Time [F (3.025, 57.48) = 19.48, p<0.0001]; Treatment [F (3, 19) = 27.08, p<0.0001]; Time × Treatment [F (12, 76) = 7.417, p<0.0001]). Similar to the females, the male mice injected with CFA developed thermal hypersensitivity 24 hours later as revealed by a Bonferroni’s post-hoc test (Fig. 2A: Day 1, Control + Saline vs CFA + Saline or CFA + MS15203, p<0.0001). The Bonferroni’s post-hoc analysis also revealed that acute MS15203 did not improve thermal latencies in CFA-treated male mice (Fig. 2A: Day 1 post-drug, Control + Saline vs CFA + MS15203, p<0.01). However, 3 days of once-daily MS15203 treatment increased thermal latencies of male mice and this improvement was sustained through the 5-day chronic treatment paradigm (Fig. 2A: Day 3 post-drug, CFA + Saline vs CFA + MS15203, p<0.05 and Day 5 post-drug, CFA + Saline vs CFA + MS15203, p<0.001).

**Fig. 2.**
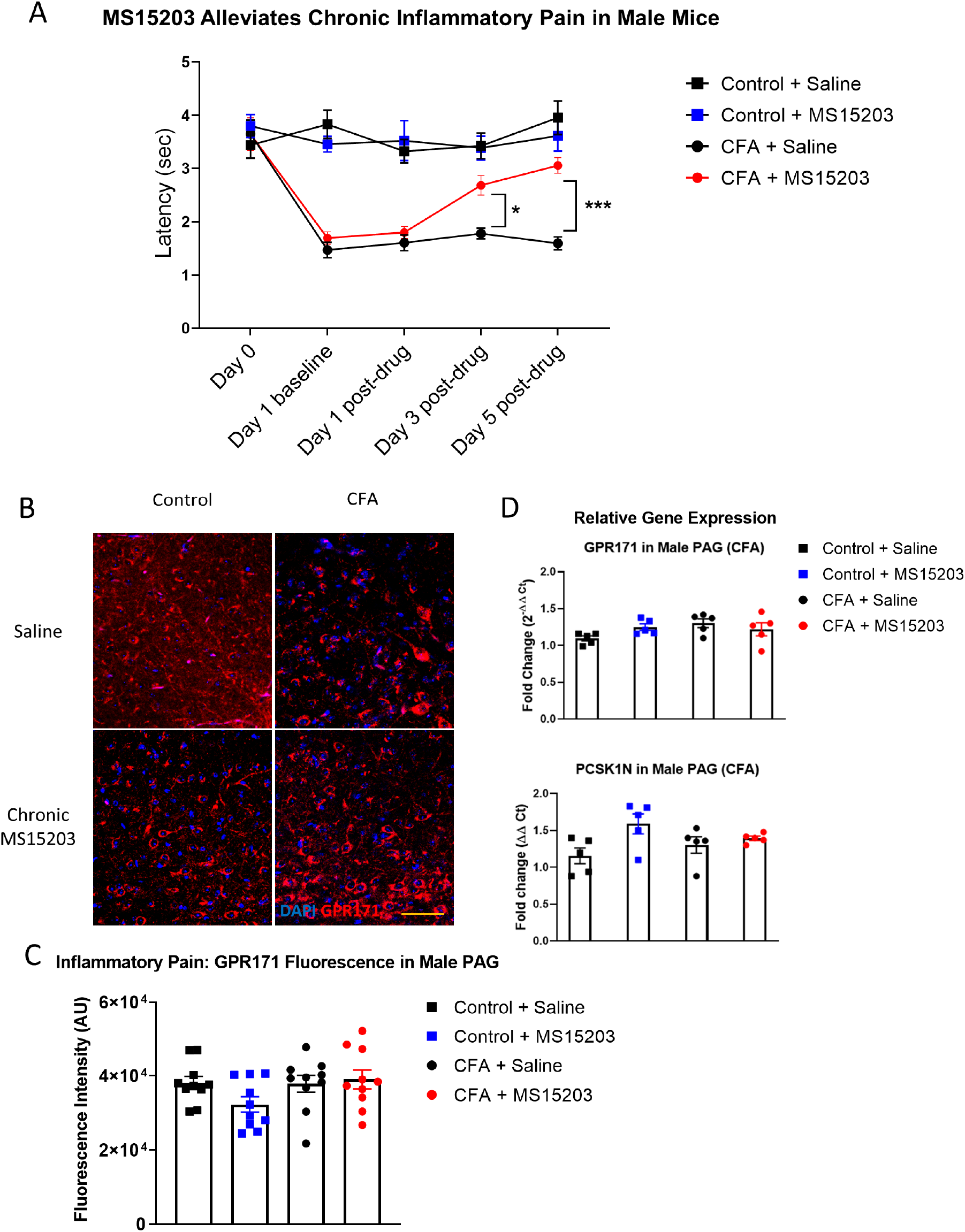
GPR171 agonist reduces chronic inflammatory pain in male mice. (A) Male mice (n=5-6/group) injected with CFA in their hind paws developed rapid thermal hypersensitivity as measured by a plantar test. Chronic treatment with MS15203 reduced the duration of inflammatory pain by Day 3 of the treatment and the analgesia was ongoing through Day 5 as measured by an increase in thermal latencies on the plantar test. Repeated measured two-way ANOVA with Bonferroni’s post-hoc test. (B) Fluorescence immunostaining and (C) quantification of GPR171 signal show that the protein levels of GPR171 are unchanged in the ventrolateral PAG (vlPAG) of male mice (n=10-12 images each from 2 mice/group). (D) Analysis of gene expression changes within the PAG reveals that the transcript levels of GPR171 and its endogenous ligand, PCSK1N, are unchanged in male mice (n=4-5/group). One-way ANOVA with Tukey’s post-hoc test. ****p<0.0001. Scale bar = 50 *µ*m.

We performed immunofluorescence staining and quantification of GPR171 in the PAG of male mice to visualize receptor localization and evaluate GPR171 protein levels within the ventrolateral PAG (vlPAG) (Fig. 2: C, D). A one-way ANOVA of fluorescence intensities indicated no significant differences between treatment groups on quantification of GPR171 protein levels in the vlPAG (Treatment F (3, 36) = 1.97, p>0.05).

We then performed qRT-PCR on dissected whole PAG from males to evaluate gene expression changes in GPR171 and PCSK1N (Fig. 2B). A one-way ANOVA indicated no significant differences between treatment groups on evaluation of GPR171 (Treatment F (3, 16) = 2.06, p>0.05) or PCSK1N (Treatment F (3, 16) = 3.07, p>0.05) expression in the PAG.

### C. GPR171 agonist does not reduce chronic neuropathic pain in female mice

We employed a model of chemotherapy-induced peripheral neuropathy (CIPN) to study the effects of GPR171 agonist treatment on chronic pain (Fig. 3A). In females (n=6-7/group), paclitaxel produced allodynia as assessed by an electronic von Frey sensor. The estrus stage did not impact the subjects’ mechanical thresholds (Fig. S1B). A repeated measures two-way ANOVA indicated significant differences in mechanical thresholds over the course of the study (Time [F (3.77, 98) = 63.87, p<0.0001]; Treatment [F (3, 26) = 44.36, p<0.0001]; Time x Treatment [F (15, 130) = 12.54, p<0.0001]). A Bonferroni’s post-hoc test revealed that the female mice developed allodynia by Day 5 following the first dose of paclitaxel (Fig. 3B: Day-5, Vehicle + Saline vs Paclitaxel + Saline or Paclitaxel + MS15203, p<0.001). Allodynia was maintained through the duration of the study and was ongoing on Day 15 prior to the initiation of pharmacological intervention (Fig. 3B, Day - 15, Vehicle + Saline vs. Paclitaxel + Saline or Paclitaxel + MS15203, p<0.0001). We then tested the effect of MS15203 at both acute (30min) and chronic (5 days) dosing paradigms. A Bonferroni’s post-hoc analysis revealed that neither acute nor chronic MS15203 treatment increased the mechanical thresholds of Paclitaxel + MS15203-treated females (Fig. 3B: Day 15 post-test and Day 20, Vehicle + Saline vs Paclitaxel + MS15203, p<0.0001). The compound MS15203 alone did not have any effect in vehicle-treated female mice.

**Fig. 3.**
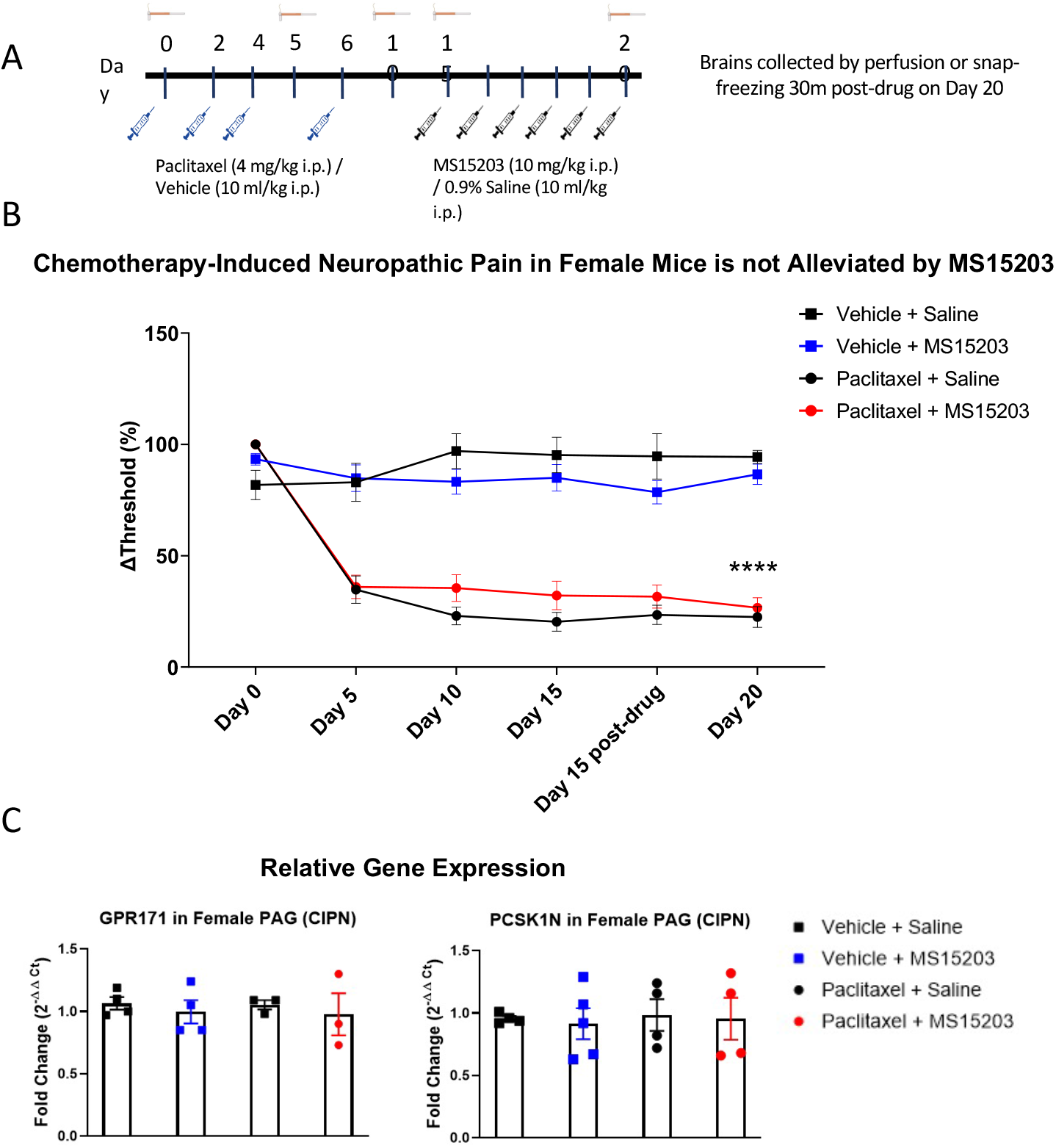
GPR171 agonist does not reduce chronic neuropathic pain in female mice. (A) An illustrative timeline describing the experimental design of chemotherapy-induced peripheral neuropathy followed by MS15203 treatment. (B) Female mice (n=6-7/group) treated with paclitaxel (16 mg/kg cumulative, i.p.) developed allodynia by Day 5 of the study as measured by the von Frey test. Chronic treatment with MS15203 from Day 15 through Day 20 failed to alleviate allodynia as noted by the persistence of mechanical hypersensitivity through Day 20. Repeated measures two-way ANOVA with Bonferroni’s post-hoc test. (C) Analysis of gene expression changes within the periaqueductal gray (PAG) reveals that the transcript levels of GPR171 and its endogenous ligand, PCSK1N, are unchanged in female mice (n=4-5/group) irrespective of neuropathic pain status or MS15203 treatment. One-way ANOVA with Tukey’s post-hoc test.

We proceeded to perform qRT-PCR on dissected PAG from female mice (4-5/group) to assess whether gene expression levels of GPR171 and its endogenous ligand PCSK1N are altered following neuropathic pain and MS15203 treatment. A one-way ANOVA revealed that the expression levels of GPR171 are unchanged in the PAG of female mice irrespective of pain condition or MS15203 treatment (F (3,13) = 0.572, p>0.05). Similarly, the expression levels of PCSK1N were also unchanged in the PAG of female mice across pain conditions or MS15203 treatment (F (3,13) = 0.076, p>0.05).

### D. GPR171 agonist reduces chronic neuropathic pain in male mice

We performed a similar experimental paradigm as Fig. 3A using male mice to assess the anti-allodynic effect of MS15203. In males (n=6-7/group), paclitaxel produced allodynia as assessed by manual von Frey filaments using the up-down method. A repeated measures two-way ANOVA indicated significant differences in mechanical thresholds over the course of the study (Time [F (3.31, 92.71) = 38, p<0.0001]; Treatment [F (3, 28) = 82.53, p<0.0001]; Time x Treatment [F (15, 140) = 13.36, p<0.0001]). A Bonferroni’s post-hoc test revealed that the male mice developed allodynia by Day 5 following the first dose of paclitaxel (Fig. 4A: Day-5, Vehicle + Saline vs Paclitaxel + Saline or Paclitaxel + MS15203, p<0.001). Allodynia was maintained through the duration of the study and was ongoing on Day 15 prior to the initiation of pharmacological intervention (Fig. 4A, Day - 15, Vehicle + Saline vs Paclitaxel + Saline or Paclitaxel + MS15203, p<0.0001). We then tested the effect of MS15203 at both acute (30min) and chronic (5 days) dosing paradigms. A Bonferroni’s post-hoc analysis revealed that acute MS15203 treatment did not alter the mechanical thresholds of male mice in neuropathic pain (Fig. 4A: Paclitaxel + MS15203 vs Paclitaxel + Saline, Day 15 baseline vs post-test, Bonferroni p>0.05). However, following 5 days of repeated dosing, MS15203 treatment increased the mechanical thresholds of mice in neuropathic pain compared to their saline-treated counterparts (Fig. 4A: Paclitaxel + MS15203 vs Paclitaxel + Saline, Day 20, Bonferroni p<0.05). The compound MS15203 alone did not have any effect in vehicle-treated male mice.

**Fig. 4.**
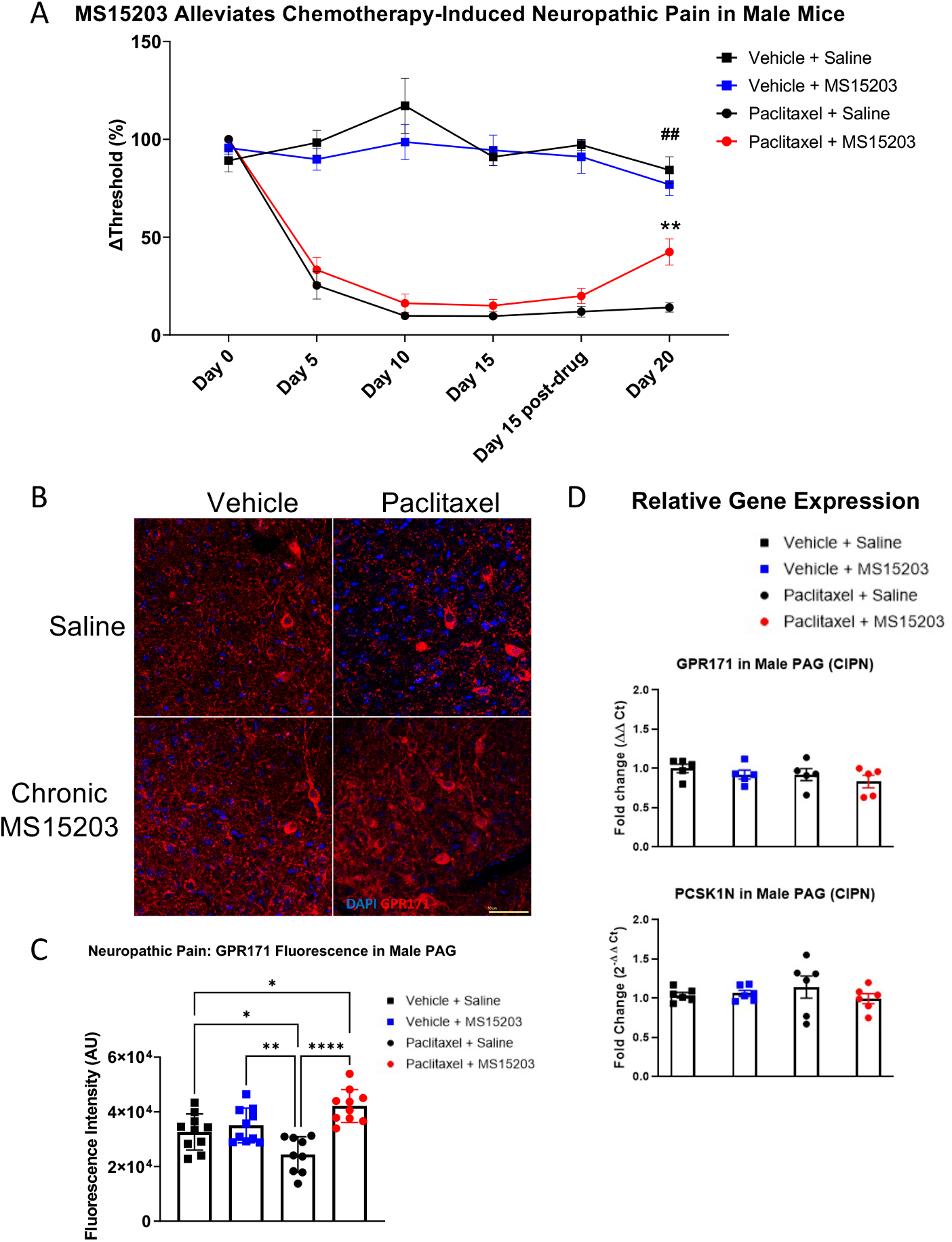
GPR171 agonist reduces chronic neuropathic pain in male mice. (A) Male mice (n=6-7/group) treated with paclitaxel (16 mg/kg cumulative, i.p.) developed allodynia by Day 5 of the study as measured by the von Frey test. Chronic, but not acute, treatment with MS15203 reduced allodynia as noted by the significant increase in mechanical thresholds compared to Paclitaxel + Saline treated mice. Repeated measures two-way ANOVA with Bonferroni’s post-hoc test. (B) Fluorescence immunostaining and (C) quantification of GPR171 signal show that protein levels of GPR171 are decreased following neuropathic pain and chronic treatment with MS15203 results in an increase of GPR171 in the vlPAG (10-12 images each from 2 mice/group). (D) Analysis of gene expression changes within the PAG reveals that the transcript levels of GPR171 and its endogenous ligand, PCSK1N, are unchanged in male mice (n=4-5/group) irrespective of neuropathic pain status or MS15203 treatment. One-way ANOVA with Tukey’s post-hoc test. *p<0.05, **p<0.01, ****p<0.0001. Scale bar = 50 *µ*m.

We performed immunostaining for the receptor GPR171 and quantified fluorescence intensity within the ventrolateral PAG (vlPAG) (Fig. 4B,C). Intriguingly, a one-way ANOVA of the fluorescence intensities revealed significant differences between the groups (F (3,35) = 10.15, p<0.0001). A Tukey’s post-hoc test revealed that neuropathic pain resulted in a significant decrease of GPR171 immunostaining compared to vehicle-treated controls (Fig. 4D: Vehicle + Saline vs Paclitaxel + Saline, Tukey’s p<0.05). Further, the post-hoc analysis revealed that following chronic MS15203 treatment, there was a rescue effect and, in fact, the levels of GPR171 were elevated compared to the vehicle-treated controls (Fig. 4D: Vehicle + Saline vs Paclitaxel + MS15203, Tukey’s p<0.05).

We proceeded to perform qRT-PCR on dissected PAG from male mice to assess whether gene expression levels of GPR171 and PCSK1N are altered following neuropathic pain and MS15203 (Fig. 4D). A one-way ANOVA revealed that the expression levels of GPR171 are unchanged in the PAG of male mice irrespective of pain condition or MS15203 treatment (F (3,13) = 0.572, p>0.05). Similarly, the expression levels of PCSK1N were also unchanged in the PAG of male mice across pain conditions or MS15203 treatment (F (3,13) = 0.076, p>0.05).

## Discussion

The current study establishes that the receptor GPR171 is a promising target for the treatment of chronic pain in males. A synthetic agonist for the receptor, MS15203, decreases the duration of both allodynia caused by neuropathic pain and thermal hypersensitivity caused by inflammatory pain in male mice. Interestingly, MS15203 does not reduce allodynia nor thermal pain in female mice using the same dose administered in males. Further, although GPR171 receptor immunostaining is unaltered in males after chronic inflammation, neuropathic pain induces a decrease in GPR171 protein levels which is rescued following chronic MS15203 treatment. The gene expression levels of GPR171 and its endogenous ligand, PCSK1, are unaltered in the PAG. Further, while GPR171 activation is recognized to promote food intake (37), we note that our 5-day treatment did not result in significant alterations in the subjects’ weights (Fig. S2A-D).

The development of CFA-induced inflammatory pain occurs over a biphasic response time. The initial course of inflammation occurs over the first 24 hours, followed by persistent pain lasting over the course of a week. We report here that systemic administration of the compound MS15203 decreases the duration of CFA-induced inflammatory pain, in a sex-dependent manner, after 3 days of treatment following the initial inflammatory phase. We also note that while this reduction of chronic pain is sustained over the course of the study in male mice, female mice do not display any reduction of thermal hypersensitivity over the course of the study. In previous studies assessing therapeutic options for chemotherapy-induced neuropathic pain, we noted that mechanical allodynia is most pronounced by Day 15 of the study and persists through 30 days (26). We assessed allodynia following 5 days of once-daily MS15203 treatment (10 mg/kg i.p.). We report here that systemic administration of the compound MS15203 decreases the duration of paclitaxel-induced peripheral neuropathy and associated allodynia after 5 days of treatment in male mice. While the pathophysiology of sex differences in neuropathic pain induced by chemotherapy is unclear, it is indeed a remarkable observation that the GPR171 activation did not promote alleviation of allodynia in females in chronic neuropathic pain.

The absence of changes in ProSAAS or GPR171 mRNA indicate that a 5-day, once-daily treatment regimen does not alter receptor or ligand gene expression in the brain although the drug does exert a physiological effect at this timescale. A recent study showed that peripheral sensory neurons contribute to the pain relief seen with GPR171 activation (38). While this observation explains alterations at the peripheral level in addition to spinal sensitization to chronic pain, the role of alterations in PAG connectivity and excitability is well established in the context of chronic pain paradigms (39–41). The absence of gene expression changes of GPR171 or PCSK1N within the PAG, while observing a modulation of protein levels is indeed a remarkable observation. We postulate that the transcript levels obtained from whole dissected PAG are not representative of the protein levels within the local vlPAG region, a highly variable relationship that has been reviewed previously (42). The decrease in GPR171 protein levels in the vlPAG following neuropathic pain is comparable to reports of decreased mu opioid receptor availability following chronic pain (43). We previously found synergistic antinociceptive effects of MS15203 with morphine, indicating the receptors could also be similar in their physiological modulation.

GPR171 is an inhibitory G*α*_i/o_ coupled receptor which inhibits cAMP production (37). Based on our study showing GPR171 in GABA neurons in the PAG, we hypothesize that activation of GPR171 leads to a decrease in GABA release (20). This in turn leads to antinociception by excitation of output neurons in the medulla. In addition, GPR171 is found in the dorsal root ganglion and alleviates inflammatory pain following intrathecal administration of a GPR171 agonist, presumably by inhibiting TRP ion channels (38). Since we administered the agonist systemically, it is likely that it is working within the central and peripheral nervous system to alleviate pain. In addition, the increase in the endogenous ligand ProSAAS, seen in circulating CSF of fibromyalgia patients, is likely a consequence of adaptations towards restoring GPR171 signaling as we have observed a decrease in brain-specific GPR171 receptor expression in males with neuropathic pain (18).

The lack of alleviation of neuropathic or inflammatory pain in females following chronic MS15203 treatment warrants further investigation. Previous studies have indicated that C57BL/6 mice display stable pain behaviors across estrus stages (44, 45). It is plausible that the dose of agonist administered (10mg/kg, i.p.) was insufficient to produce a physiological response in females. Further, there is considerable evidence of heightened immune activation in females at both central and peripheral sensory processing regions, which contribute to mechanical and thermal hypersensitivity (11, 46–48). The heightened microglial activation in the PAG in females has been shown to reduce morphine-induced antinociception (49). While there is a paucity of studies examining immune interactions with GPR171 activity, the possibility of the immune microenvironment modulating GPR171-dependent analgesic activity cannot be ruled out. Further, studies with female mice have reported that the sex-specific differences in chronic pain are driven in part by the action of hormones and are peripherally regulated by TRP channels (44, 50). As GPR171 exerts its peripheral effects via TRP channels on sensory afferents in male mice, the microenvironment in this region can be postulated to contribute to the observed sex differences (38). Further, as the gene encoding GPR171 is located on chromosome 3 of the mouse, the observed differences in behavior cannot be attributed to sex chromosome-linked causes. Indeed, sex differences in antinociception have been found in other systems such as the fatty acid-derived resolvin D5, that was able to produce antinociception in males with neuropathic or inflammatory pain, but not in females (51). Our findings thus identify GPR171 not only as a novel target for the treatment of chronic inflammatory and neuropathic pain, but also one that displays sexual dimorphism in pain regulation.

Chronic inflammatory and neuropathic pain present public health concerns without efficacious treatment options. Our findings suggest that the receptor GPR171 is a therapeutic target for the treatment of multiple modalities of chronic pain in a sex-dependent manner and its agonist, MS15203, can be used to treat chronic pain.

## ACKNOWLEDGMENTS

This study was supported by startup funds from the Department of Biology at Utah State University, a Pharmacology and Toxicology startup grant from the PhRMA Foundation and Young Investigator Grant from the Brain and Behavior Research Foundation (to ENB). The authors thank Dr. Mona Buhusi and Valerie Martin (Utah State University) for timely assistance during the COVID-19 pandemic with supplies essential for the completion of the study. The authors thank Dr. Sanjai Pathak (Queen’s College, New York) for the kind gift of MS15203.

## Supplementary Material

**Fig. S1.**
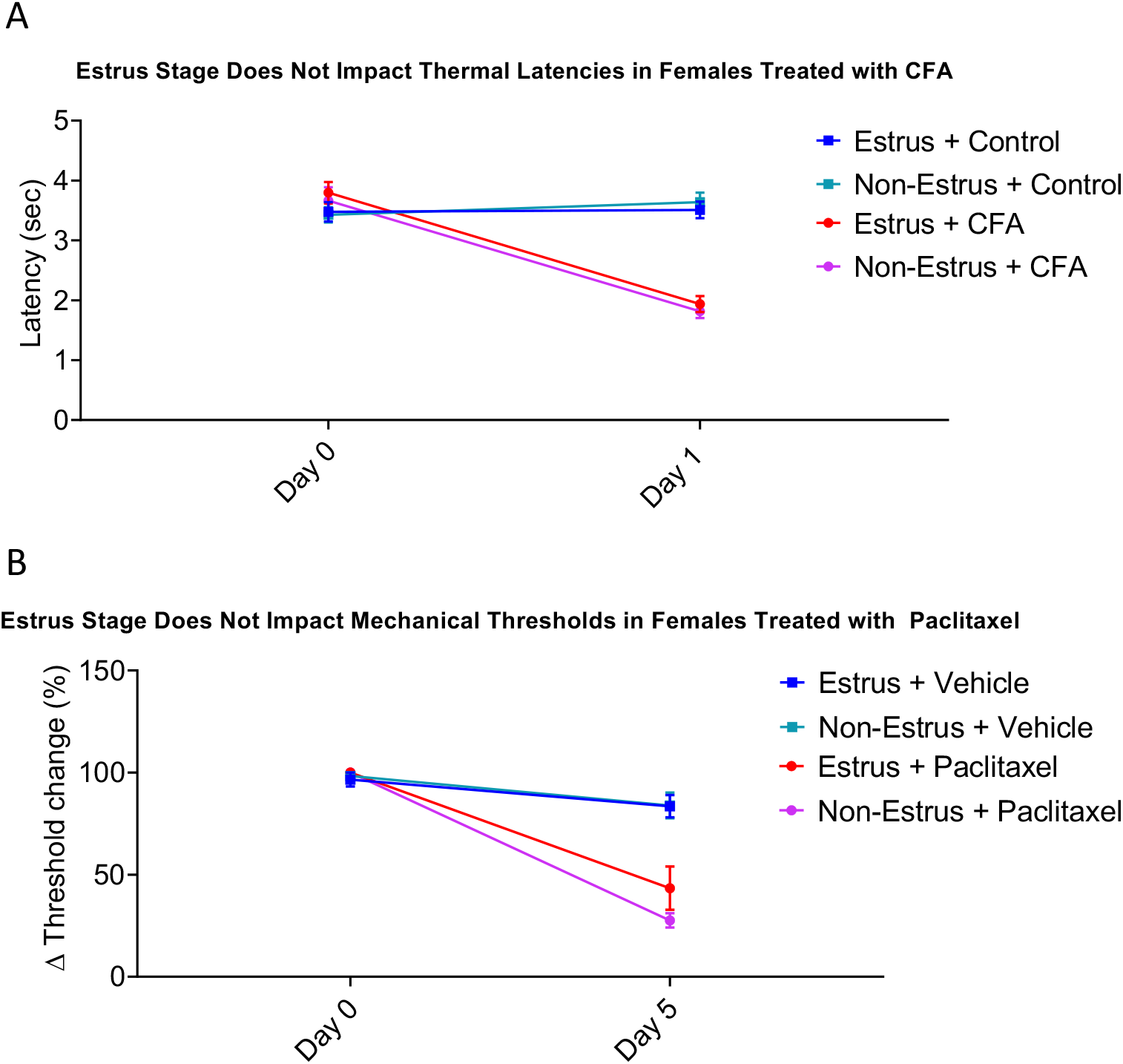
Estrus stage does not impact baseline thresholds of female mice. Female mice do not show significant differences in their A) thermal thresholds or B) mechanical thresholds on Day 0 of the chronic pain studies. Two-way repeated measures ANOVA indicated significant differences in the thresholds of the mice in inflammatory or neuropathic pain. However, a Bonferroni’s post-hoc test revealed that there were no differences in the baseline thresholds of mice in their respective studies.

**Table S1.**
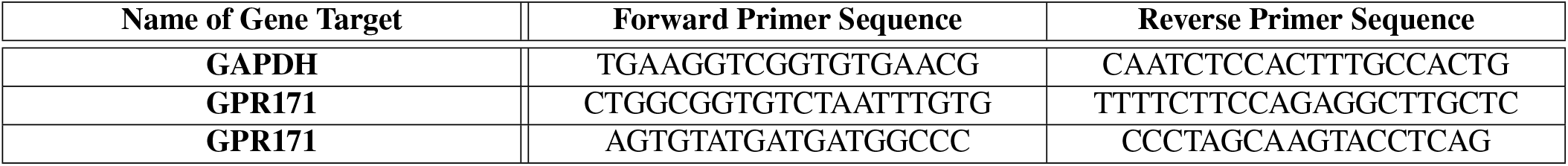
Primer sequences for quantitative RT-PCR.

**Fig. S2.**
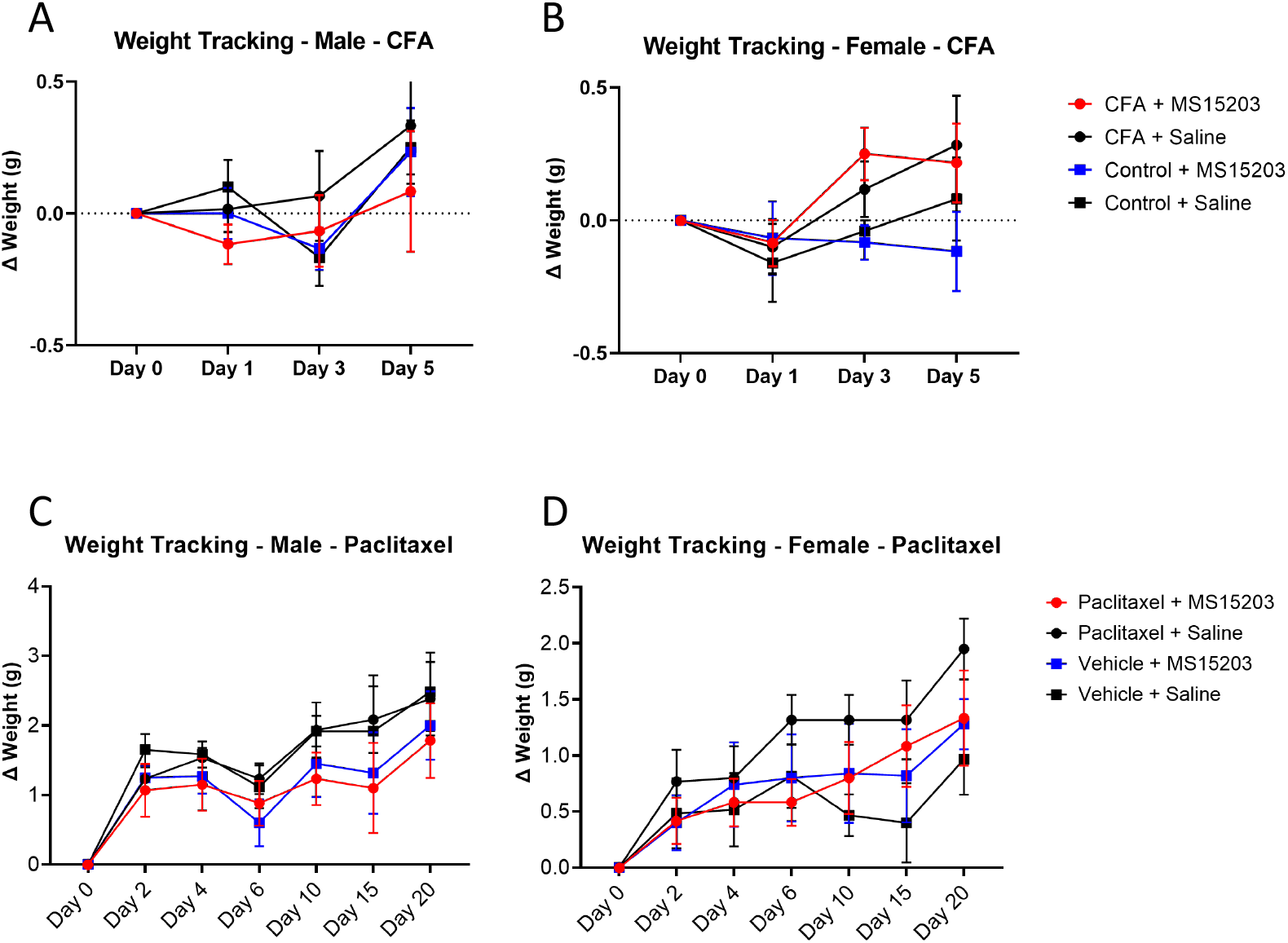
MS15203 treatment does not produce significant weight gain in male or female mice. Once-daily injection of MS15203 (10mg/kg, i.p.) for 5 days does not produce significant weight grain in male (A, C) or female (B, D) mice. A two-way repeated measures ANOVA indicated significant differences in weight changes in mice in the chemotherapy-induced peripheral neuropathy (CIPN) study. However, a Bonferroni’s post-hoc test revealed that there were no significant differences in weight changes between Day 15 and Day 20 of the study, the duration in which the mice received MS15203.

